# Investigating face and house discrimination at foveal to parafoveal locations reveals category-specific characteristics

**DOI:** 10.1101/807099

**Authors:** Olga Kreichman, Yoram Bonneh, Sharon Gilaie-Dotan

## Abstract

Since perceptual and neural face sensitivity is associated with a foveal bias, and neural place sensitivity is associated with a peripheral bias (integration over space), we hypothesized that face perception ability will decline more with eccentricity than place perception ability. We also hypothesized that face perception ability may show an upper visual field (UVF) bias due to the proximity of face-related regions to UVF retinotopic representations, and a left visual field (LeVF) bias due to earlier reports suggesting right hemisphere dominance for faces. Participants performed foveal and parafoveal face and house discrimination tasks for upright or inverted stimuli (≤ 4°) while their eye movements were monitored. Low-level visual tasks were also measured. The eccentricity-related accuracy reductions were evident for all categories. Through detailed analyses we found (i) a robust face inversion effect across the parafovea, while for houses an opposite effect was found, (ii) higher eccentricity-related sensitivity for face performance than for house performance (via inverted vs. upright within-category eccentricity-driven reductions), (iii) within-category but not across-category performance associations across eccentricities, and (iv) no hemifield biases. Our central to parafoveal investigations suggest that high-level vision processing may be reflected in behavioural performance.

## Introduction

One of the main coding principles in the visual cortex is eccentricity, where the foveal representation is significantly magnified in the visual cortex relative to its size on the retina, and as distance from the fovea grows, the cortical representation is reduced ^1–3^. This foveal magnification is assumed to be the main factor contributing to reduced performance with growing eccentricity for multiple (but not all) visual tasks ^4–6^. Studies also show that this foveal enhancement can be overcome if peripheral information is scaled to match the cortical representation (aka M-scaling ^7^).

Another coding principle is hierarchical processing, where visual information is initially processed according to the physical aspects in early visual areas (e.g. in V1, V2) and as information progresses in the visual hierarchy it is integrated and becomes associated with perceptual aspects (specialization) rather than the physical aspects of the stimulus in higher visual areas (e.g. in LOC, FFA, PPA)^8–14^. This high-level visual specialization is predominantly manifested in two anatomically distinct visual processing pathways, the ventral perception visual pathway processing aspects related to shape and form, and the dorsal spatial visual pathway processing aspects related to action preparation and location in space ^15–19^.

Multiple neuroimaging and electrophysiological studies investigating the ventral perception visual pathway have revealed that face processing is associated with foveally biased regions and critically rely on them while place and house related processing are associated with and rely on peripherally biased regions ^20–30^ sensitive to integration over space ^20,31^. In addition, population receptive field (pRF) studies with faces show that most of the pRFs in the ventral face areas include the foveal area, and extend to cover parafoveal locations during peripheral face task ^32,33^. Recent studies further suggest that the structure of these category selective regions is also distinct ^25,26^.

Reasoning that neural coding preferences likely reflect behavioural performance, and based on earlier studies and on reduced visual performance at peripheral locations ^6^, we hypothesized that face related tasks, that are associated with a cortical foveal bias would show reduced performance in more peripheral locations relative to house related tasks that are associated with a cortical peripheral bias. Thus we expected that place related tasks would be less affected by eccentricity and would show, on top of the expected eccentricity reductions ^6^, higher performance than faces as eccentricity increases (as presented in the middle and left models in Fig. 1a). While we would like to compare face and house related performance, accuracy levels may be affected by across category task difficulty differences. To account for potential across categories performance differences at the center (Fig. 1a on the left) each category’s performance can be normalized according to central performance (Fig. 1a middle panel). Such normalization allows examining the effects of the eccentricity-related performance reductions relative to central performance. However, we cannot rule out the possibility that despite differences in neural representations, face and house performance will not show significant differences across foveal to parafoveal locations (see Fig. 1a right model). Furthermore, we also hypothesized that due to the proximity of the ventral face-related regions in the fusiform to upper visual field (UVF) retinotopic representations (at the posterior ventral aspects of the visual cortex (e.g.^34,35^), face perception ability may show an UVF bias ^36^ or a left visual field (LeVF) bias due to earlier reports suggesting right hemisphere dominance for faces^37–44^. Lastly, due to the evident VF biases in scene/place-related regions in pRF studies (UVF bias in ventral PPA and LoVF bias in dorsal TOS ^45^), we also hypothesized that there may be an UVF or LoVF difference in house performance. To test these hypotheses we measured face and house discrimination performance for upright and inverted stimuli at foveal to parafoveal locations (up to 4°, see Figure 1b-e) in each of the four visual field quadrants in a group of normally sighted individuals while participants held fixation and their eye movements were being monitored. We reasoned that since inversion has been used to expose holistic processing of central faces (e.g. ^46–49^), measuring how eccentricity modulates inversion may expose characteristics reflecting category-specific mechanisms. To minimize potential task difficulty related differences between face and house performance, on top of examining performance in the same visual field locations, we also examined normalized performances, reductions in performance levels as a function of eccentricity (i.e. slopes), and how inversion is modulated by eccentricity. In addition we reasoned that if category-related performance relies on category-specific mechanisms we should expect that despite the eccentricity-related reductions in performance (i) performance across eccentricities would be correlated for a specific category, but (ii) performance across categories should not be correlated.

**Figure 1.**
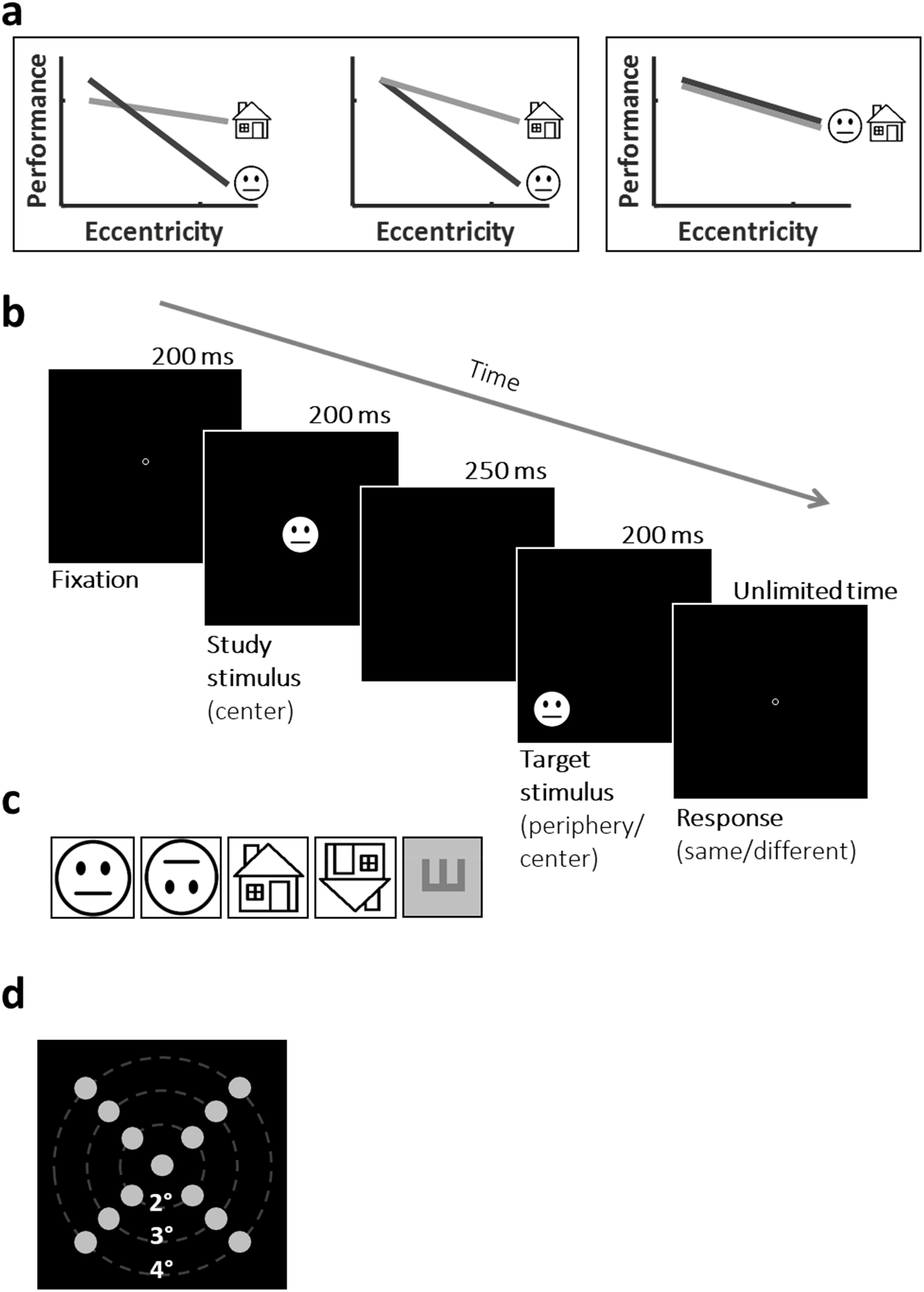
Predicted outcomes and experimental design. (a) Proposed models for face and place-related processing as would be reflected in face and house discrimination performance across the parafoveal eccentricities. Left and middle panels – both house and face discrimination are affected by eccentricity but to a different extent. Due to possible between-categories task difficulty differences, we cannot compare central performance of houses and faces (e.g. left panel) and therefore the middle panel presents hypothesized normalized performance for each category. These panels predict that house discrimination would be less affected by eccentricity than faces, i.e. showing less reduction in performance across parafoveal eccentricities. Right-face discrimination would not show superiority to house discrimination at any parafoveal location. (b) Representative timeline of a face discrimination “same” trial at 4° eccentricity (left lower visual field). Each trial started with a fixation circle appearing for 200ms followed by a central study face appearing for 200ms, and after a 250ms ISI a target face appeared for 200ms in 1 of 13 randomly chosen locations at central or parafoveal eccentricities (see panel d). The participant’s task was to report if the target face was the same as (“same” condition) or different than (“different” condition) the study face. (c) Representative stimuli from each of the 4 different category discrimination experiments. (d) The 13 possible locations for the target stimuli in the category discrimination experiments (see panel c). In each discrimination experiment there were 50 trials in each location (25 “same”, and 25 “different” trials). In the E discrimination experiment we only tested performance at central, 2° and 4° eccentricity locations, overall 9 locations with 40 trials in each location (20 “same”, and 20 “different’ trials). Note that the experimental stimuli were real faces and of real houses and are fully described in the Methods section and the ones depicted here are only for illustration.

## Results

In line with the eccentricity expected reductions in performance, we found that for all visual categories accuracy and d-prime were highest at the centre of the visual field and dropped significantly as eccentricity increased (see Figure 2, Table 1 for detailed results, and Table 2 for statistical results). The superior performance at the centre of the visual field was also evident in longer RTs for growing eccentricities for all categories and orientations (see Figure 2c-d, Table 1 for detailed results, Table 2 for statistical results, and Supp. Mat.).

**Figure 2.**
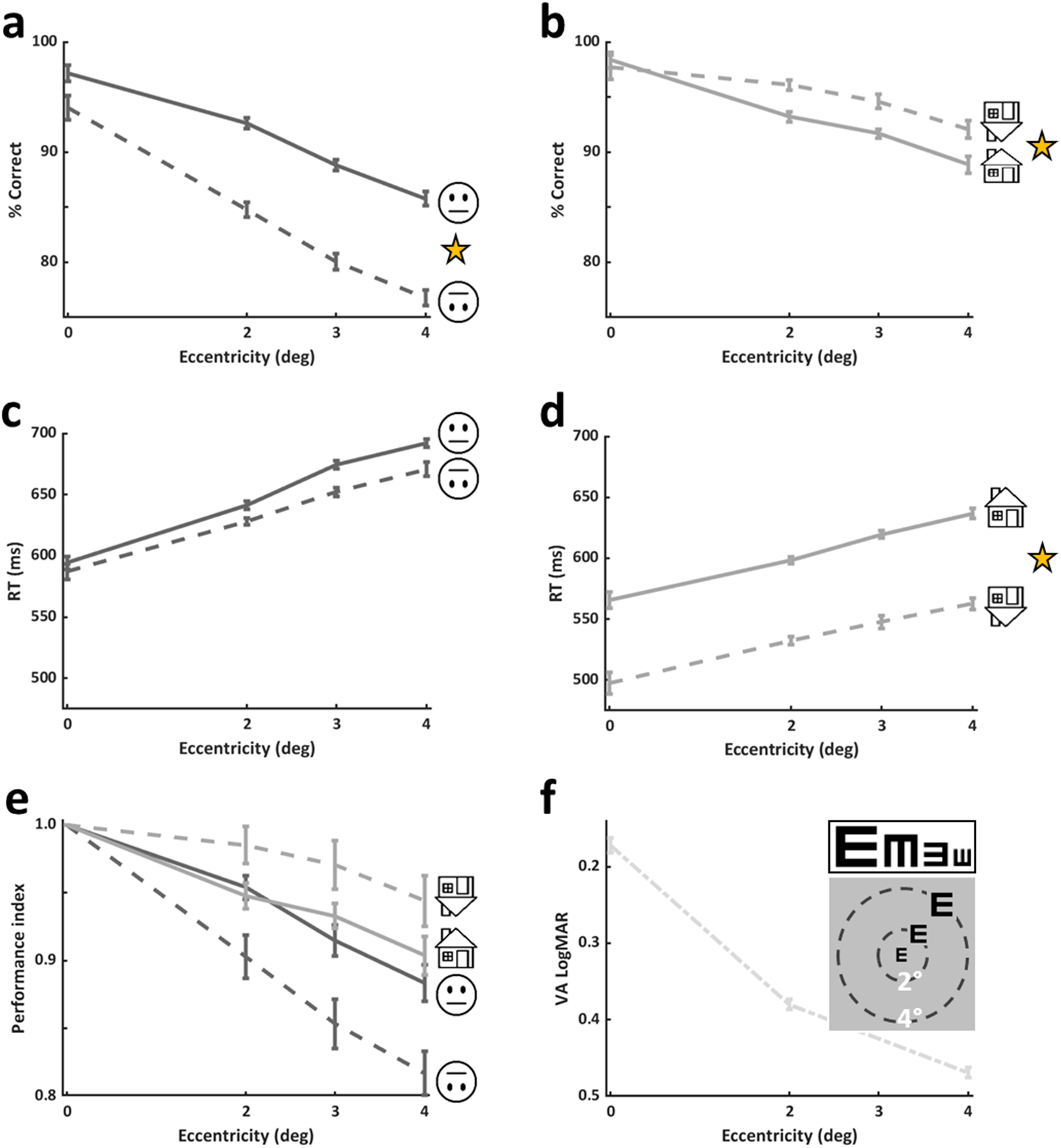
Experimental results. Upright and inverted accuracy for faces (a) and for houses (b). As can be seen performance declines with eccentricity for faces and for houses (see Results). There was a significant face inversion effect across parafoveal eccentricities (main effect of inversion in a 2-way ANOVA on accuracies with eccentricity and orientation: F(1,196)=72.2, p<0.0001) evident in reduced performance for inverted faces relative to upright ones. A significant house inversion effect in the opposite direction to the face inversion effect was found (main effect of inversion in a 2-way ANOVA on accuracies with eccentricity and orientation: F(1,140)=7.8, p=0.005; see Results for more details). RTs (median) as a function of eccentricity for upright and inverted faces (c) and for upright and inverted houses (d). Note that RTs of inverted faces were comparable to those of upright faces, while RTs of inverted faces were significantly faster than those of upright houses (main effect of inversion in a 2-way ANOVA on RTs with eccentricity and orientation F(1,140)=27.87, p<0.0001). (e) Performance index (relative to performance at the centre of the visual field) calculated for each discrimination experiment) to estimate the rate of decline in performance relative to best vision at the centre. (f) ‘VA’ tumbling E experimental results by eccentricity. As can be seen at 2° visual acuity seems to decline relative to the centre by an average of 0.2 LogMAR units which corresponds to 2 lines on the ETDRS chart, and an additional decline of another line on the ETDRS chart for 4°. Asterisks in (a), (b), and (d) denote significant within category inversion effects. Data presented in this figure is based on upright faces (n=29), inverted faces (n=22), upright houses (n=22), and inverted houses (n=15). Error bars in (a), (b), (c), (d), and (f) represent SE calculated using the Cousineau method, error bars in (e) represent standard error of the mean (SEM). See also Supplementary Figure SF3.

**Table 1.** Summary of experimental results for each experiment by eccentricity. Mean ± SE are provided for accuracy, and performance index; median ± SE are provided for reaction times for all experiments (dprime and shape discrimination results are available in Supplementary Table ST1). Accuracy (discrimination experiments) is reported in % correct, visual acuity (tumbling E experiment) is reported in LogMAR units. Reaction time (RT) is reported in ms. Performance index represents proportion out of 1. NA - data not available. Slopes were calculated as the slope from 4° to 0° based on performance index (normalized) results. Slope difference represents the differences between upright minus inverted slopes for each category. SE for accuracy and RT is calculated based on the Cousineau method^59^, and for the other data is based on the typical formula.

|  |  | Faces |  | Houses |  | VA |
| --- | --- | --- | --- | --- | --- | --- |
|  |  | Upright | Inverted | Upright | Inverted | "Tumbling E" |
|  |  | (n=29) | (n=22) | (n=22) | (n=15) | (n=19) |
| Accuracy | Centre | 97.17 ± 0.73 | 94.03 ± 1.10 | 98.34 ± 0.74 | 97.70 ± 1.09 | 0.17 ± 0.009 |
|  | 2° | 92.63 ± 0.48 | 84.78 ± 0.68 | 93.20 ± 0.47 | 96.09 ± 0.44 | 0.38 ± 0.006 |
|  | 3° | 88.81 ± 0.49 | 80.06 ± 0.71 | 91.68 ± 0.39 | 94.60 ± 0.66 | NA |
|  | 4° | 85.75 ± 0.66 | 76.77 ± 0.68 | 88.85 ± 0.74 | 92.06 ± 0.78 | 0.46 ± 0.006 |
| RT (ms) | Centre | 594.32 ± 5.18 | 586.954 ± 6.40 | 565.79 ± 6.65 | 497.46 ± 8.71 | 568.21 ± 12.3 |
|  | 2° | 641.22 ± 3.38 | 628.15 ± 2.87 | 598.40 ± 2.87 | 632.23 ± 3.44 | 585.26 ± 8.56 |
|  | 3° | 674.46 ± 3.26 | 652.09 ± 3.741 | 619.50 ± 2.99 | 547.83 ± 5.24 | NA |
|  | 4° | 692.06 ± 3.51 | 670.65 ± 5.91 | 636.68 ± 4.25 | 562.80 ± 4.73 | 609.55 ± 8.49 |
| Performance index | Centre | 1 ± 0 | 1 ± 0 | 1 ± 0 | 1 ± 0 | NA |
|  | 2° | 0.95 ± 0.008 | 0.90 ± 0.016 | 0.94 ± 0.009 | 0.98 ± 0.013 | NA |
|  | 3° | 0.91 ± 0.011 | 0.85 ± 0.018 | 0.93 ± 0.009 | 0.97 ± 0.018 | NA |
|  | 4° | 0.88 ± 0.013 | 0.81 ± 0.016 | 0.90 ± 0.014 | 0.94 ± 0.018 | NA |
|  |  | (n=15) | (n=15) | (n=15) | (n=15) | NA |
| Slope |  | -0.029 ± 0.004 | -0.045 ± 0.005 | -0.019 ± 0.003 | -0.014 ± 0.004 | NA |
| Slope difference |  | 0.016 ± 0.005 |  | -0.005 ± 0.004 |  | NA |

**Table 2.**
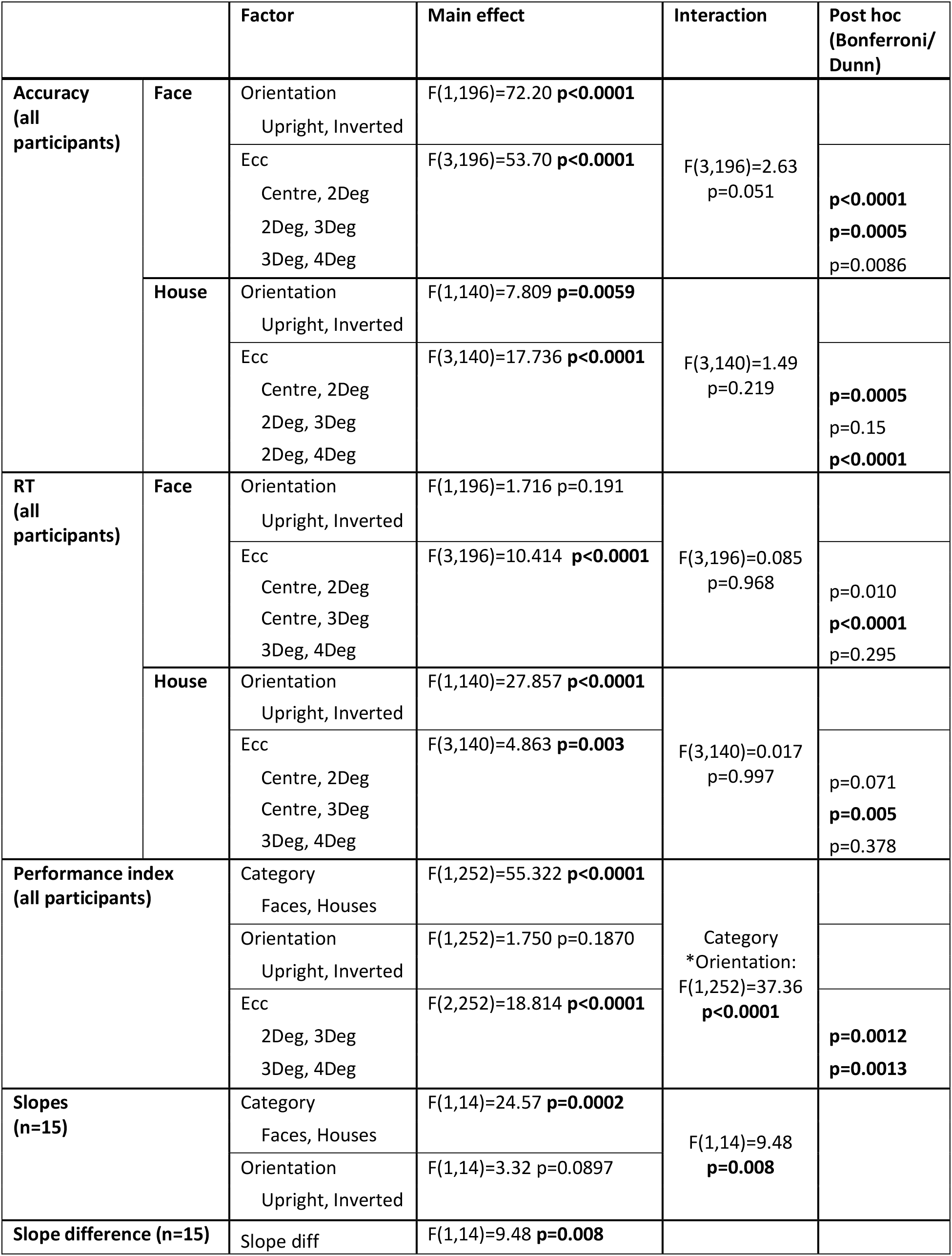
Summary of statistical analyses across experiments. Statistical analyses were performed using nonrepeated measures two-way ANOVA with eccentricity and category on accuracy or RT, 3-way non-repeated measures ANOVA for performance index (eccentricity, orientation and category), and 2-way repeated measures ANOVA (category and orientation) for slopes. This was followed by Bonferroni/Dunn post-hoc tests. F statistics are provided with corresponding p values. Values in bold represent significant results (in post-hoc surviving Bonferroni/Dunn corrections). Note that we did not find an inversion effect for houses at 0° (direct comparison of upright houses accuracy and inverted houses accuracy for all participants: parametric unequal variance t-test t(22.15)= 0.57, p=0.57, non-parametric two-sided Wilcoxon rank sum test p=0.971 z=0.036).

On top of the expected eccentricity-related reductions we found (see Figure 2 and Table 1), we directly assessed whether faces and houses are qualitatively different in a series of analyses. First, in Figure 2a and b we can see the comparison between upright and inverted performance for faces (Figure 2a) and for houses (Figure 2b). While performance was significantly higher for upright than for inverted faces (see statistical results of a 2-way ANOVA in Table 2) corresponding to the face inversion effect, for houses, following a 2-way ANOVA analysis, we found that inverted houses’ performance was significantly higher than that of upright houses (see Table 2), and for both categories this was not a result of speed-accuracy trade-off (see Figure 2 and compare panels c with a and d with b). Second, since performance as a function of eccentricity may be relative to performance at the center, we normalized accuracy measurements according to performance at the center that served as a baseline, and this was done for each participant and each category, creating performance indices. These results can be seen in Figure 2e, reflecting the opposite effects that inversion had on face (reduction) and on house (increase) discrimination (see Table 1). It can also be seen in Figure 2c and d that reaction times were differently affected by inversion for each category (see Table 1 and for statistical results see Table 2). The next set of analyses focused on a group of participants (n=15) that participated in the 4 main experiments (upright/inverted faces/houses) allowing us to perform within-participant and between-category analyses (Figures 3 and 4). Since we wanted to estimate how performance of each category is modulated by eccentricity, and since the reductions in performance from central vision to parafovea were apparently linear, we calculated for each participant the slope of the eccentricity-driven performance decline for upright and inverted faces and houses. As can be seen in Figure 3a that presents the average eccentricity-driven slopes for each category and each orientation, we found a significant effect of category and an interaction between category and stimulus orientation, also reflecting the results in Figure 2 (see Table 2 for statistical details), and this indicates that inversion affects performance differently for faces than for houses. Figure 3b displays individuals’ slopes for upright and inverted faces (dark gray) and houses (light gray). Since we hypothesized that the differences between the upright and inverted slopes may be a possible category distinguishing feature, we calculated a single characteristic measure for each category that took into account both the upright and the inverted performance modulation by eccentricity and their relations. This measure was a subtraction of the inverted slope from the upright slope for each category (see Figure 3c). This measure was found to be significantly different between the two categories (see Table 2). Individual data of this analysis are presented in Figure 3d. Lastly, following the logic of Haxby et al. 2001 ^50^, we reasoned that if faces and houses are supported by distinct category-specific mechanisms, then performance would be associated across eccentricities for each category but not across categories. On the other hand, if performance is only driven by eccentricity and not by category-specific mechanisms, then we should expect that performance across categories would be correlated. We found a significant correlation between performance index in 3° and in 4° for faces (R^2^=0.607, p=0.0006, non-directional, Fig. 4a) and for houses (R^2^=0.470, p=0.004, non-directional, Fig. 4b) supporting category specific mechanisms. Furthermore, when comparing faces at 4° and house performance at 3° (R^2^=0.095, p=0.262, non-directional, Fig. 4c) or face performance at 3° and house performance at 4° (R^2^=0.089, p=0.278, non-directional, Fig. 4d) we found that these were not significantly correlated (see Figure 4c, d) which further supports category-specific mechanisms.

**Figure 3.**
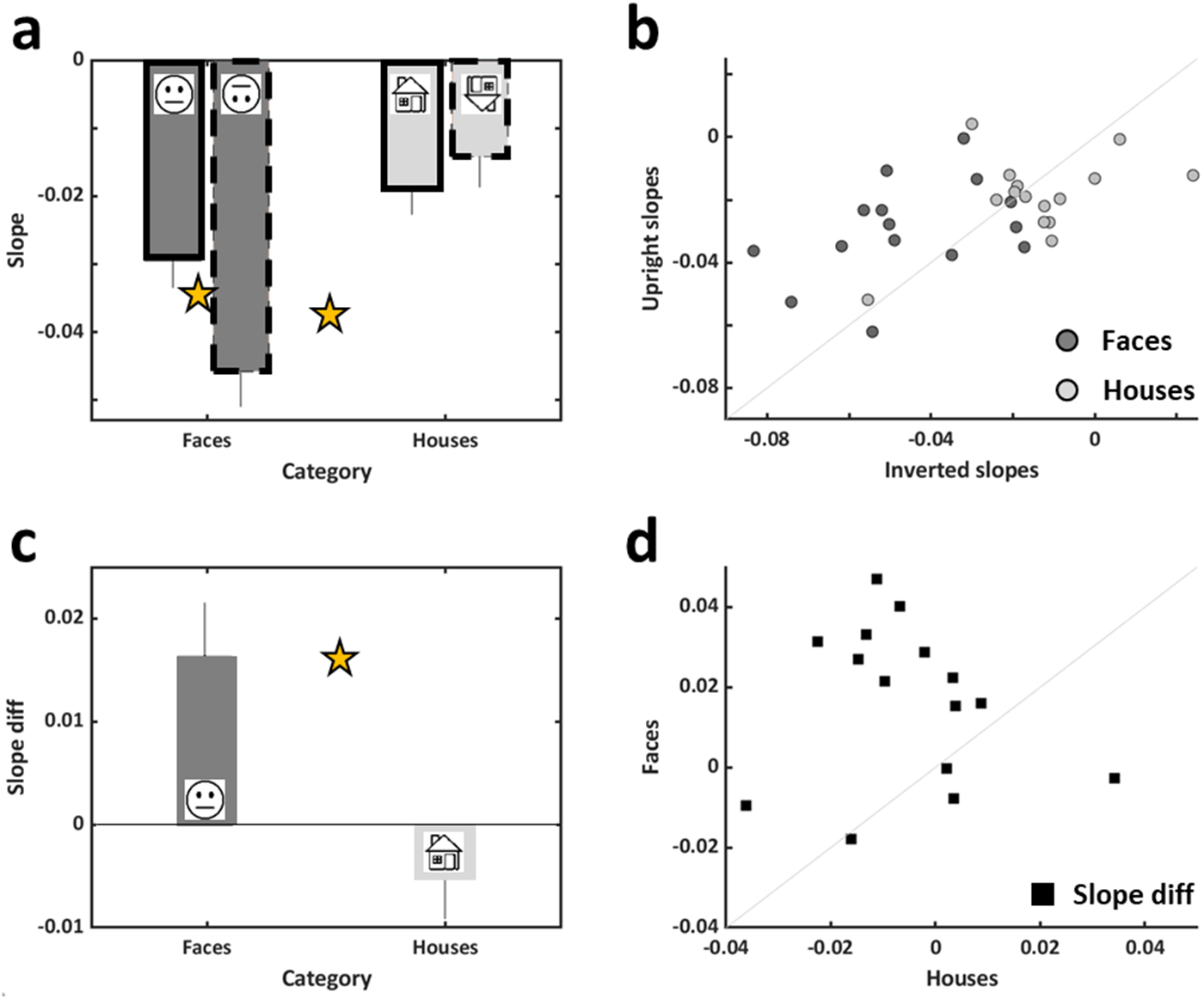
Slope analyses: Eccentricity related performance modulations (n=15). (a) The slope of the approximately linear eccentricity-driven performance decline for each category and orientation averaged across participants. The negative values correspond to the expected eccentricity effect. Significant effect of category (F(1,14)=24.57,p=0.0002, indicated by the between-category asterisk) and an interaction between category and orientation (F(1,14)=9.48, p=0.008) were found. Inverted faces’ slope was significantly bigger in magnitude than that of upright faces (p=0.006, t(14) = 3.214, 2-tailed paired t-test, marked by an asterisk), while no such difference was found for houses (p>0.23, t(14)=1.23). (b) For each participant individual slopes of upright (y axis) are plotted against inverted (x axis) ones, faces in dark grey, houses in light grey. (c) Average differences between upright and inverted slopes for faces and for houses. Each bar represents the average across all participants. A significant difference between faces and houses was found (t(14)=3.07, 0=0.008, 2-tailed paired t-test, denoted by an asterisk). See Results and Discussion for more details. (d) Individual face slope differences plotted against house slope differences. Error bars in (a) and (c) represent standard error of the mean (SEM).

**Figure 4.**
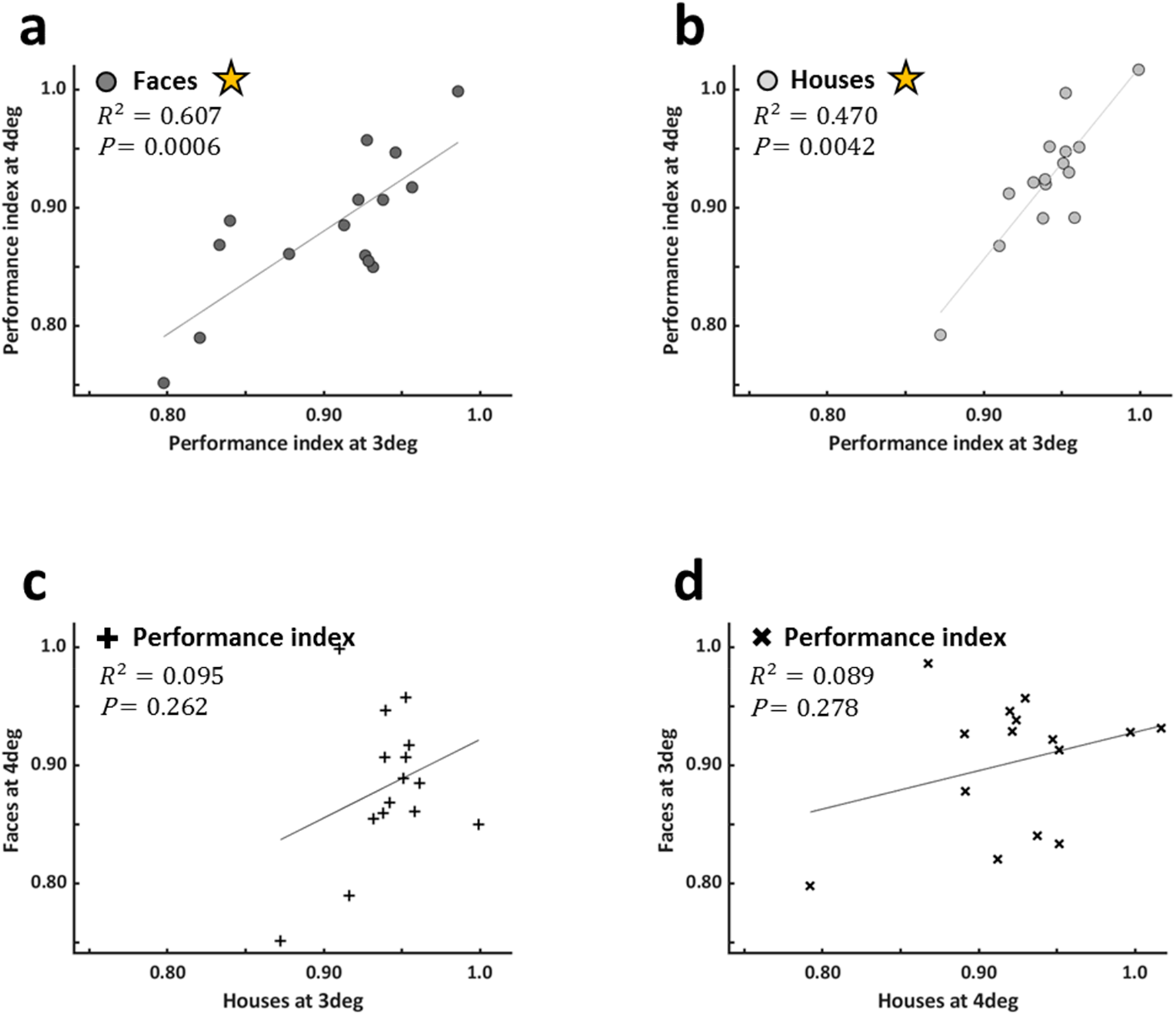
Across eccentricity within-category and across-category correlations (n=15). (a) Face performance index (PI) at 3° (x axis) is significantly correlated with face PI at 4° (y axis) (R^2^=0.607, p=0.0006, non-directional), and (b) house PI at 3° (x axis) is significantly correlated with house PI at 4° (y axis) (R^2^=0.470, p=0.004, non-directional). However, across category PIs were not significantly correlated: (c) house PI at 3 ° vs face PI at 4°, (d) face PI at 3° vs house PI at 4°. Asterisks denote significant correlations.

### Estimating visual acuity

VA related measurements were also estimated by the ‘VA’ tumbling E experiment at different locations to provide an estimate for basic vision performance decline with eccentricity. As can be seen in Figure 2f, at 2° visual acuity seemed to decline relative to the centre by an average of 0.2 LogMAR units which correspond to 2 lines on the ETDRS chart, and an additional decline of another line on the ETDRS chart for 4°. Typically, a value of 0 LogMAR corresponds to static VA of 6/6, and a 0.1 difference reflects one line in the static chart.

### Visual field comparisons

As can be seen in Figure 5 and in Supplementary Tables ST2 and ST3, no significant differences between upper visual field (UVF, in blue) and lower visual field (LoVF, in green) or between left visual field (LeVF, in red) and right visual field (RVF, in yellow) were found for upright faces or for any other categories.

**Figure 5.**
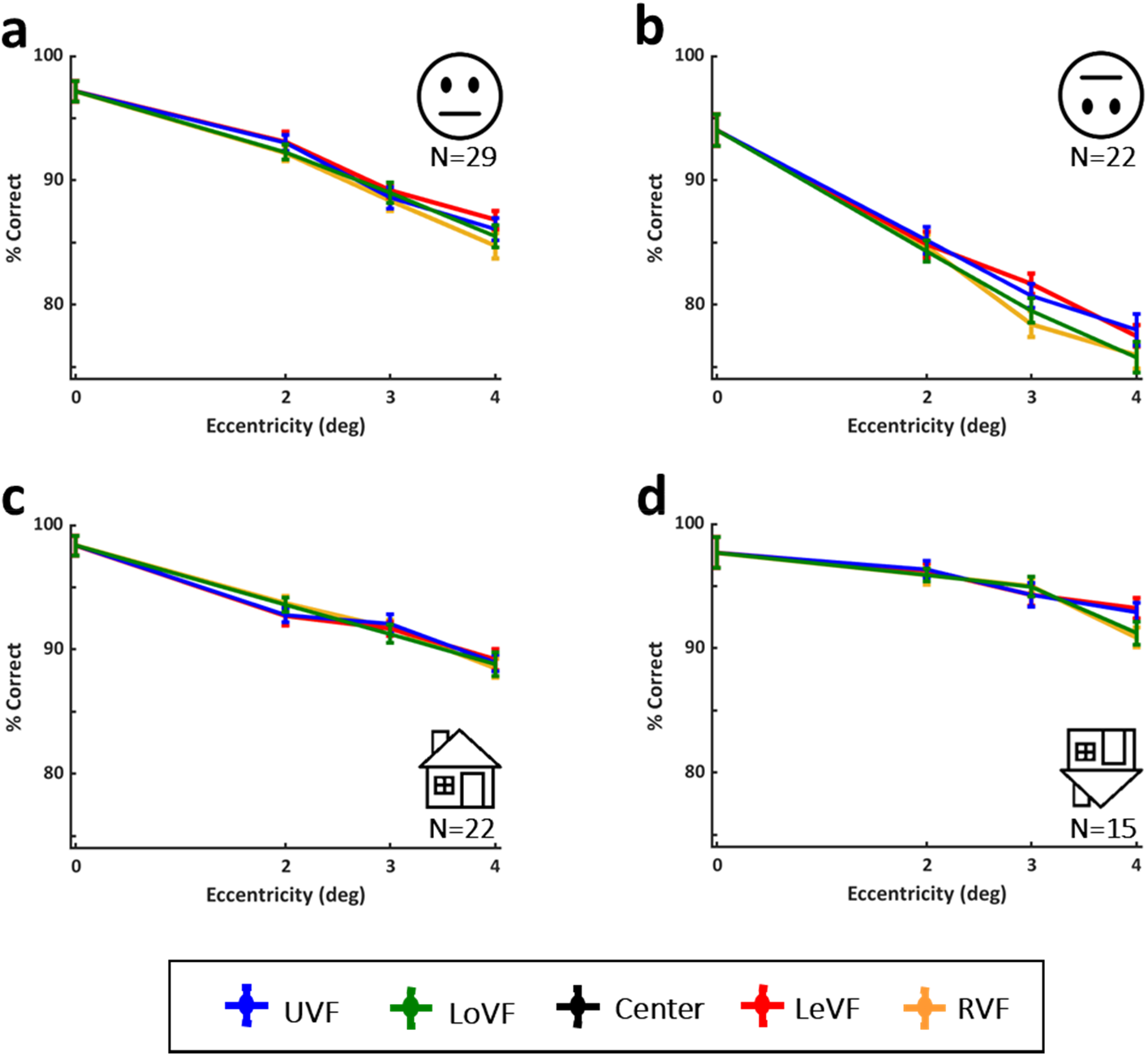
Accuracy for the upright and inverted faces and houses discrimination experiments by visual fields. Performance in each of the discrimination tasks ((a) upright faces (n=29), (b) inverted faces (n=22), (c) upright houses (n=22), (d) inverted houses (n=15)) declines with eccentricity. No significant differences between upper (in blue) and lower (in green) visual fields or between right (in yellow) and left (in red) visual fields were found for any category. Error bars represent standard error calculated by the Cousineau method.

### Estimating physical differences’ contribution to performance

We calculated the physical differences between sets of images used in our experiments to examine whether these could possibly account for performance. To that end, we estimated for each pair of different images used in the discrimination house or face experiments the physical difference between them (Euclidean distance) and examined whether bigger physical differences would be associated with higher performance (see Methods for more details). We reasoned that a bigger physical difference would facilitate distinguishing between different images (and thus improving accuracy on the ‘different’ condition). Figure 6 shows the physical difference between pairs of images for each experiment (Fig. 6a), accuracy performance (Fig. 6b) and RTs (Fig. 6C) for faces and houses for the different house discrimination experimental versions (see details in Methods) as a function of eccentricity. As can be seen, accuracy of house version 1 was higher than that of house version 2 (post-hoc p=0.004), but house version 2 was not significantly different than face accuracy (post-hoc p=0.241, see also Supplementary Tables ST4 for details and ST5 for statistical results). However, despite the significant physical differences between the house versions, there was no significant difference between the RTs of the two house versions (see ST5). The physical difference between the house versions can explain the differences in accuracy between the house versions (version 1 with higher physical differences between the images may have made house discrimination easier, and indeed showed higher accuracy than version 2 with the smaller physical differences). However, physical differences were not likely to account for the face vs house performance differences since despite significant different physical differences between the face and house version 2 (p<0.0001), there was no significant difference between face and houses version 2 performance (p=0.241, see also Supplementary Table ST5).

**Figure 6.**
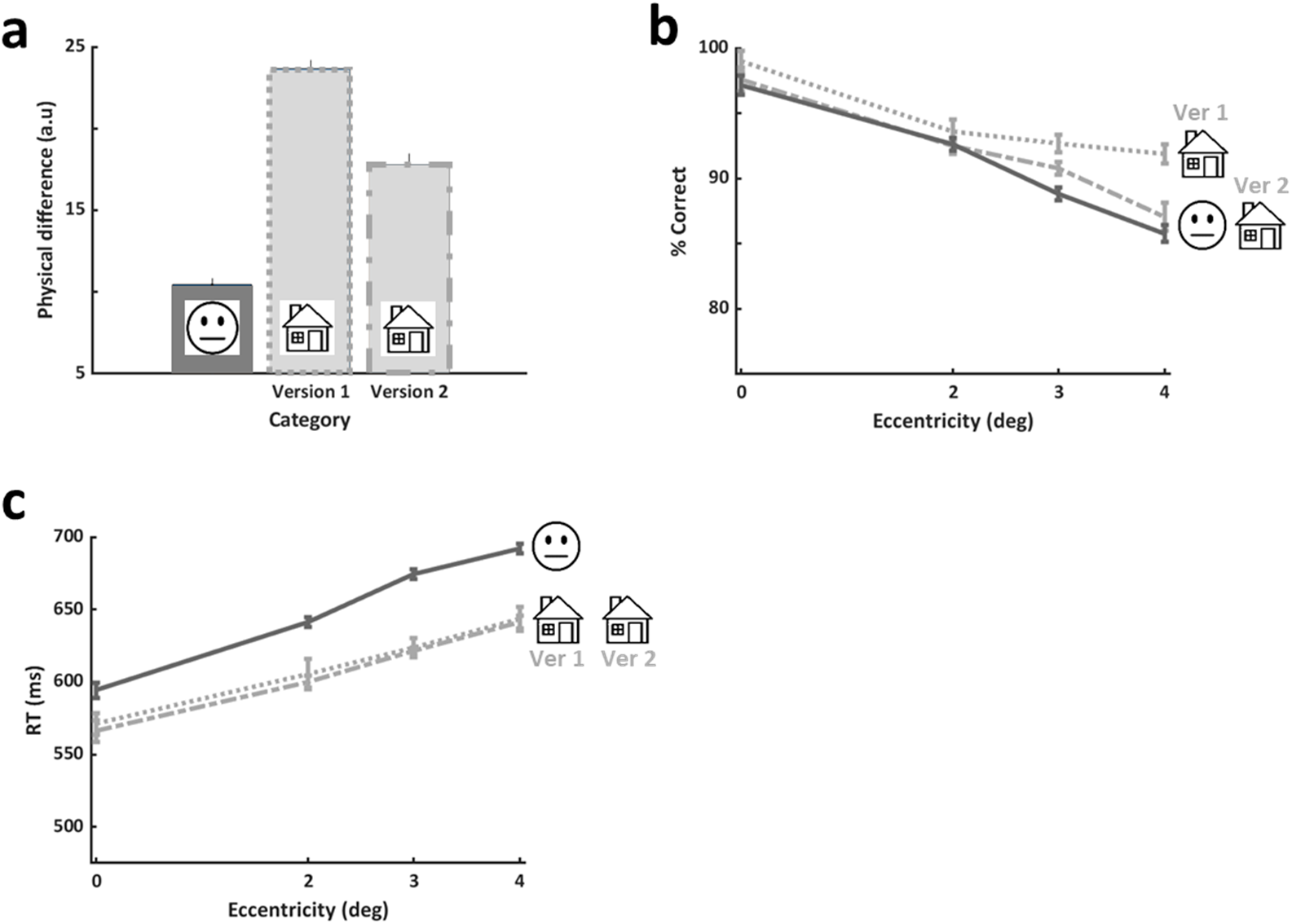
Relating low level physical differences to performance (all study participants). The physical differences between image pairs (either from the upright face experiment, or from each of the upright house discrimination versions) were compared to performance differences. (a) Mean physical difference between pairs of stimuli in the face discrimination task (on the left) and two versions of the house discrimination task (Ver1 in the middle and Ver2 on the right). There was a significant difference between the two house versions, and house Ver1 had bigger physical difference than that of faces (one way ANOVA on physical difference between experiments: main effect of experiment (F(2,72)=143.42, p<0.0001, post-hoc Bonferroni/Dunn Ver1-Ver2: p<0.0001, Ver2-Faces: p<0.0001). (b) Accuracy by eccentricity for the different experimental versions. A significant difference in performance between the house discrimination versions was found with no difference between the face experiment and house Ver2 (2-way non-repeated measures ANOVA on accuracy with experiment (house Ver1, house Ver2, Faces) and eccentricity revealed a main effect of version: F(2,280)=9.23, p=0.0001; post-hoc Bonferroni/Dunn Ver1-Ver2: p=0.004, Ver2-Faces: p>0.24). (c) RTs by eccentricity for the different experimental versions. No significant difference between the RTs of both house discrimination versions were found, but as reported in the Results, houses RTs were significantly different (faster) than those of face discrimination (post-hoc Bonferroni/Dunn: Ver1-faces: p=0.003, Ver2-faces: p=0.001). Note that accuracy differences may have reflected within category differences, and RTs may have reflected between category physical differences. Nevertheless, inversion, which did not change a category’s physical differences, significantly affected within category performance for both faces and for houses (see Results). Therefore, physical differences may only partially account for performance. Error bars in (a) represent standard error of the mean (SEM), error bars in (b) and (c) represent standard errors calculated using the Cousineau method.

## Methods

### Participants

A group of 41 neurotypical participants aged 18-47 (mean age 26.6 years ± 6 (SD)) with normal or corrected-to-normal vision participated in this study. After exclusion (see criteria below) 29 participants took part in the upright face discrimination experiment, 22 of them participated in the inverted face discrimination experiment, 18 of them and 4 additional participants (altogether 22 participants) participated in the upright house discrimination experiment, 15 of them participated in the inverted house discrimination experiment, 18 of them and an additional participant (altogether 19 participants) participated in the tumbling E experiment^51^, and 6 of them and additional 4 participants (altogether 10 participants) participated in the shape discrimination experiment. All participants signed an informed consent form before their participation. The Bar Ilan University ethics committee approved all the experimental protocol, and the methods were carried out according to the ethics committee guidelines and regulations. Participants whose accuracy or RTs exceeded 2SD from the mean of the group in any of the experiments were excluded from all the study analyses.

### Apparatus

All experiments were conducted on an Eizo FG2421 24” HD LCD monitor (1920×1080 pixels resolution) running at 100 Hz, using an in-house developed platform for psychophysical and eye-tracking experiments (PSY) developed by Yoram S. Bonneh ^52^ running on a Windows PC.

### Eye tracking

In order to ensure fixation during all experimental sessions, eye movements were recorded with an EyeLink infrared system (SR Research, Ontario, Canada) with a sampling rate of 500Hz, equipped with a 35mm lens while head movements were limited by chin rest. Eye movements were recorded binocularly, only left eye data were analyzed. A standard 5-point calibration was performed before each session. In addition, in-session calibration trials were incorporated into the beginning and end of each session to estimate the deviation from fixation for each session. This was computed as the difference between the eye position during the in-session position calibration (baseline) and the average eye position during that session. Trials with a deviation from fixation greater than 0.5° during a time window around the onset of the target stimulus were excluded from the analysis.

### General procedure

Participants were instructed to look at the fixation point and be alert to and aware of the surrounding area in which the target could appear. Each experimental session included eye position calibration trials before and after the main experiment (see more details below). Throughout all experimental sessions participants started each trial at their own pace by pressing a key after fixation appeared at the centre of the screen. Experiments were performed in a dark room, no feedback was given, and the viewing distance was 60 cm.

### Main experiments

#### 1. Upright faces discrimination experiment

In each trial, participants were asked to judge whether two faces presented sequentially were the same or different. A study face always appeared at the centre of the screen for 200ms, and after an ISI of 250ms, a target face (either same or different than the study face) appeared for 200ms (to eliminate the possibility of succeeding in the task if performing a saccade towards the target) at a location chosen randomly from 13 possible locations in the visual field (centre or one of 4 locations at 2°, 3° or at 4°, as depicted in Figure 1d). The face images for the same condition (same image served as the study and the target face) were chosen randomly from a set of 5 different faces, each appeared once in each location per session. For the different condition, each of the two face images was chosen randomly from two different sets of 5 face images each, 5 of the face images used for the different condition were the images used in the same condition. All face images were full-front colored photographs of real men with a neutral expression cropped and aligned to each other (see full details at ^53,54^ with the original images taken from 2 databases (CVL Face Database [http://www.lrv.fri.uni-lj.si/facedb.html]; AR Face Database [Martinez and Benavente 1998]). The faces were presented on a black background and subtended 1.6° x 2.2° (width x height). Trials were mixed randomly in terms of condition (same/different) and location in the visual field. Each participant underwent 5 runs of the experiment with each condition (same/different) repeated 5 times in each of the 13 locations. Overall, there were 25 ‘same’ face trials and 25 ‘different’ face trials for each of the 13 locations for each participant. Face discrimination performance was measured as accuracy (percent correct) per location. Experimental procedures are illustrated in Figure 1b-d.

#### 2. Upright houses discrimination experiment

The experimental design was identical to that used in the face discrimination paradigm except for the use of grayscale images of real houses rather than the face images. We ran two versions of the experiment, each using a different set of house images. The first version was built to equate central performance between faces and houses. However, since the physical difference between the house and face experiments was big, we created an additional house version with a different set of houses to reduce the physical between stimuli differences. Each participant performed the first version (V1) twice and the second version (V2) 3 times; order of sessions was V1-V2-V2-V1-V2. House images subtended 2.4° x 2° (width x height). Experimental procedures are illustrated in Figure 1b-d.

#### 3. Inverted faces discrimination experiment

The experimental design was identical to that used in the face discrimination paradigm except that faces were inverted (both study and target faces). Experimental procedures are illustrated in Figure 1b-e.

#### 4. Inverted houses discrimination experiment

The experimental design was identical to that used in the upright houses discrimination paradigm except that houses were inverted (both study and target houses). Experimental procedures are illustrated in Figure 1b-e.

#### 5. VA tumbling ‘E’ experiment

A tumbling E test was used to measure visual acuity (VA) threshold ^51^ at 9 different visual field locations (centre, 2° and 4°, 4 locations at each eccentricity). Separate staircase procedures were applied for each of the 9 locations; trials of all locations were mixed randomly in each session. The stimuli were a black E on gray background that in each location subtended initially 0.5° and faced 1 of 4 optional directions. Participants’ task was to determine the E’s facing direction (4AFC), while the E’s size was reduced in a 3:1 staircase procedure with 0.1 log unit steps according to performance (stopping after 6 reversals). Experimental procedure is illustrated in Figure 1d-e and Supplementary Figure SF1. Results are reported in log MAR units (minimum angle of resolution (MAR) needed to correctly discriminate (at 79% accuracy performance) the E’s facing direction, values closer to 0 indicate better VA).

Note that our measurements of dynamic VA are not precisely comparable to the standard static ETDRS measurements since (i) our viewing distance (60 cm) was not standard (for near 40 cm or for far 3 m), (ii) exposure time was limited (vs. unlimited exposure in static VA examinations), (iii) divided attention across the VF (vs. focusing all attention on the centre of VF), (iv) contrast was lower than used in standard VA tests. However, the dynamic tumbling E measurements have been shown to correspond to VA standard measures ^57,58^.

Shape discrimination experimental details are provided in Supplementary Material.

### Analysis

All analyses reported were calculated individually and then the average over participants and SE are reported.

For each of the discrimination experiments performance was measured as accuracy (percent correct) per location and for the VA experiment performance was reported in log MAR units per location (as presented in Figure 2). For each location we calculated individual performance and we present the average results over all participants. For each experiment and each eccentricity, we averaged the performance of all 4 locations (upper/lower left/right locations) of that eccentricity. RT analyses were done in the same way as for accuracy measures.

In order to assess the eccentricity effect (i.e. the drop in visual performance with eccentricity) and compare it across all discrimination tasks, we computed a *performance index* (*PI*), see Figure 2E. PI was calculated as the performance (accuracy) at each eccentricity divided by the performance at the centre. In the group of the n=15 that performed all 4 main experiments (upright/inverted faces and houses) we measured the eccentricity related reductions in performance by calculating the *slopes* of the decline from central vision (0 eccentricity) to 4° of eccentricity by subtracting the performance at 0° from that of 4° (the average performance across all 4 locations in that eccentricity), and these are presented in Figure 3a and b. *Slope differences* were calculated for each visual category (faces and houses) as the difference between the upright slope and the inverted slope (slope difference = upright slope – inverted slope). As before, this was calculated for each participant and then averaged across participants. Results are presented in Figure 3c and d.

To compare upper vs. lower VF and the right vs. the left VF performances, the two locations at each eccentricity and each hemifield were averaged (e.g. to compare UVF vs. LoVF, for the 2°, 2° right UVF and left UVF were averaged and compared to the average of 2° right LoVF and left LoVF), see Figure 5 and Supplementary Figure SF2.

To estimate *physical distances* (or *differences*) that may account for differences in performance, we calculated the Euclidean distance between 2 different face images (for faces) or 2 different house images (for houses). Euclidean distance between 2 images was estimated as the mean luminance level absolute difference over all pixels in these images. These are presented in Figure 6a.

Statistical analyses (ANOVA and post-hoc) were performed with StatView5.0 software for Windows (SAS Institute Inc, Cary, NC) and presented in Tables 2, Supplementary Table ST3 and ST5.

Error bars for accuracy, RTs, and VA represent standard errors across participants calculated using the Cousineau method (see ^59^ for details). For the performance indices and slope-related analyses (see below) error bars represent standard error of the mean (SEM).

## Discussion

Our investigations into face and house-related processing in central to parafoveal locations (up to 4°) revealed that, as expected, all tested categories, showed an eccentricity effect (i.e. reduced accuracy and d-prime, and slower RTs with growing eccentricity^6^). Our findings also support our hypothesis that place-related processing would be less affected by eccentricity than face related processing (as proposed in both left and middle models in Fig. 1a), based on earlier neuroimaging findings that faces are associated with foveal bias and places with a peripheral one ^20,21,23–26,29,30^. Our hypothesis was supported by larger slopes of eccentricity related reductions for faces than for houses. In addition, for each category the slope of eccentricity related reductions was modulated quantitatively (i.e. magnitude) and qualitatively (i.e. direction) different by inversion suggesting different processing for faces and for houses in the parafovea. Specifically, accuracy of inverted faces was worse than that of upright faces at all eccentricities, which extended the central vision face inversion effect ^47,60,61^ to the parafovea. In contrast, for houses, we found an opposite effect with higher performance for inverted stimuli. Furthermore, we found that within-category but not between-category performance was associated across eccentricities, again supporting dissociated category-related mechanisms which is in line with functional and structural brain studies ^25,50^. In addition, we found no significant performance biases for LeVF, RVF, upper or lower visual fields (see also Supplementary Material).

Our study aimed to assess whether high-level visual behavioural performance reflects brain organization as evident from functional and anatomical brain studies of human visual cortex. Multiple neuroimaging and electrophysiological studies ^20,23–26,30^ show that faces are associated with foveal processing and houses and places with peripheral processing and integration over space ^31^. We anticipated that this will be reflected in behaviour according to the model proposed in Figure 1a left and middle panels. In earlier neuroimaging studies that investigated eccentricity biases in face and place related regions, the goal was to determine whether the overall eccentricity biases coexisted or were evident in category sensitive regions (using category specific stimuli of up to ~6° from the fovea, and overall general visual stimuli of up to ~10° from the fovea)^20,30^. In our study we were interested to investigate how category specific behaviour is modulated by eccentricity within parafoveal domain (≤4°). Some earlier neuroimaging studies have also addressed the relation between category related areas and the effects of stimulus size on their activation levels (e.g. ^20,31,62^). Some of these studies show that the activity in face-related areas is not affected by image size changes for central stimuli while the activity in place-related areas is influenced by size. Another study shows that for peripheral stimuli (at 10°) image size does seem to enhance the face-related N170 component, and this may be due to the larger images penetrating the central receptive fields of the face areas ^33^. In our study we kept the stimulus sizes constant across the parafovea reasoning that as in natural vision, objects’ world size does not change. Under these experimental conditions and while refraining from direct between-category comparisons, our behavioural results support high level vision brain organization principles for faces and places (see above).

When comparing visual categories in brain imaging research most often less emphasis is given to controlling task difficulty across categories (e.g. ^20,21,23,50^). However, in our study we wanted to investigate whether and how the extensively studied high-level visual brain mechanisms of faces and places are reflected in behaviour. To study this behaviourally we wanted to compare between categories. Therefore, we needed to control for differences in task difficulty between the categories that despite our attempts to equate them may arise from the different processing characteristics of each category (foveal vs peripheral bias, differences in spectral features or in physical world size) or from our experimental design. To that end we refrained from directly comparing between accuracy levels of the two categories. Instead, we exploited inversion since stimulus inversion is known to adversely affect the holistic nature of face perception ^46–49^ and since both categories are typically perceived in an upright orientation. We used inversion as a category-related measure to assess within-category inversion effects across eccentricities, and these were compared between faces and places. This enabled us to relate behaviour to neuroimaging studies investigating faces and places. Our results are in line with the model suggesting that faces and places are differentially modulated by eccentricity in the parafovea as depicted in Figure 1a left and middle panels. In line with neuroimaging results, we found that faces are more adversely affected by eccentricity than houses. This was evident in a series of analyses that we performed (see Results) resulting in a single characteristic measure (slope difference, see Results and Methods) that indicated that the magnitude of modulation for faces by eccentricity is bigger than that of houses. The fact that the faces’ slope difference was in an opposite direction to that of houses (see Figure 3c), and that within category (but not across categories) performance was correlated across eccentricities support the idea that the mechanisms underlying face and place processing are dissociated (for review see ^25^), and our study extends this to parafoveal vision.

Since faces are known to show a foveal bias, it is yet unclear how face inversion will be modulated by eccentricity. Although inverted faces keep the local features and spatial relations of the faces intact, the stimuli and their holistic structure are unfamiliar to the visual system and this leads to the face inversion effect ^47,60,61^, where people are worse when performing a task on inverted faces relative to upright faces. Earlier studies show that processing of inverted faces recruit both face specific and non-face related mechanisms that include parietal attention-related foci^54,60,63–65^. An earlier study examining face inversion at right or left peripheral locations compensated for the foveal cortical magnification by enlarging peripheral faces (by CMF) reported on an inversion effect in the periphery without any RT differences ^48^. However, we were interested to investigate how the face inversion effect is modulated by eccentricity when face size is kept constant, as is typical during natural vision. One possibility is that face inversion would be evident in peripheral vision if upright and inverted faces performance would both proportionally degrade with growing eccentricity. Another possibility is that in peripheral vision upright face perception will reduce and reach levels similar to those of inverted faces, thus weakening the central face inversion effect. Our design allowed us to examine these two possibilities in both accuracies and response times. Interestingly, while we did not find any effects of inversion on RTs, we found that the face inversion effect in central vision (that was evident in accuracies) did not diminish with eccentricity, and in fact seemed to grow (see the bigger slope for inverted faces relative to that of upright ones).

While inversion is known to affect central face perception and we have also found that is extends to parafoveal vision, it is unclear how inversion affects other visual categories. Here we examined how inversion affects an additional category of houses, which is considered in the neuroimaging literature to be an opposing category to faces (e.g. ^13,14,66^). We found, in contrast to face inversion, that accuracy was higher and RTs were faster for discriminating inverted houses relative to upright houses, and this was similar to what was found earlier for central faces and houses ^67^. Thus this inversion effect was different than that found for faces. Several possible explanations may account for this. First, since the house inversion experiments in our study were performed after the upright houses experiments, one could argue that learning effects may underlie the improved performance for inverted houses. However, since this was also the case for faces (with inverted faces being always after upright ones), this explanation may not be the only factor explaining the differences between house and face inversion effects. A second explanation is related to the peripheral bias associated with places. A direct comparison of upright vs inverted central houses did not reveal any significant accuracy difference, and the significant effect we found between upright and inverted houses stems from differences in parafoveal eccentricities (see Figure 2b). In contrast to faces, mechanisms supporting houses show a periphery related effect ^20,24,31^, and therefore may be more sensitive to inversion (manipulation of stimulus orientation) at the periphery rather than in central vision. This may explain why faces show an inversion effect at the centre of the visual field which grows with eccentricity, while houses are sensitive to inversion only with peripheral stimuli which does not seem to grow with eccentricity (no significant slope difference between the slopes of upright and inverted houses). Thirdly, inversion may reduce holistic processing, and this in turn may lead to local low level feature comparisons, which are assumed to be quicker. This would clearly disrupt holistic face processing but may simplify house discrimination, and the faster RTs for inverted houses relative to upright ones may provide an indirect indication for this. We observed a different pattern of inversion effects on houses than on faces, and thus we anticipate that inversion may influence additional visual categories in a category specific manner (for a review see ^68^). It is possible that face and place mark the limits of this behavioural effect despite possibly relying on different brain mechanisms that come into play under inversion ^54,60,61,63^.

The fact that house related performance was faster than that of faces, even at the fovea, may seem counterintuitive. However, as we mentioned above, task difficulty may play a role and thus can perhaps explain why RTs for houses may be faster. Furthermore, in addition to house/place related processing relying on different cortical mechanisms and different computations than those of faces ^25,26,66,69^, house/place related mechanisms also show a tendency for transient rather than sustained activity ^66,70^. One study shows that house related areas as the house related ventral PPA and dorsal TOS areas also exhibit transient BOLD responses while the face related areas (ventral FFA and also OFA) show sustained BOLD activity and this is independent of the preferred or non-preferred category ^66^. Another recent study shows that different areas in the ventral stream receive different contributions of transient and sustained inputs ^71^. If transient related activity is associated with faster processing than that associated with sustained activity, then these differences could contribute to differences in response timing. Another possibility is that the differences in physical low level aspects of the stimulus could contribute to the speed of processing and this again may relate to task difficulty differences. As we discussed above (and see also Fig. 6a), there were bigger physical differences between pairs of house images than between pairs of face images. Thus, the house task may have been easier and thus a decision may have been reached sooner.

While we found multiple indications that face and house behaviour reflect brain mechanisms supporting face and house processing, we cannot rule out some possible confounding factors that may limit the generalization of our findings. First, our paradigm enabled us to investigate parafoveal eccentricities up to 4° and perhaps at further peripheral locations these effects may be greater or diminish for faces and for houses when peripheral mechanisms start to kick in. A second possibility relates to the small size of our stimuli with an emphasis on the house stimuli. Houses and places in our daily life are typically big and take up a bigger portion of the visual field ^27^. The small houses used in our experiment may have forced the system to rely on additional mechanisms that are non-typical to house related processes, and these non-typical mechanisms contributed to the house-related performance we found. Nevertheless most neuroimaging investigations, like our study, have been carried out with stimuli that are not directly reflecting world size. A third possibility is that low-level aspects of our specific stimulus sets could have contributed to the effects we found for faces and houses. Indeed, the analysis we carried out examining whether physical differences may account for the performances we observed (see Figure 6) indicated that physical differences may partially but not fully explain performance. Furthermore, it could be argued that all that our experimental design recruited predominantly shape discrimination mechanisms. However, if this were the case then inversion should have resulted in comparable performances. Since there were clear and significant inversion effects for faces and for houses, and they were different from each other, it is unlikely that we merely measured shape discrimination processes. A fourth possibility is that top-down attentional mechanisms influence performance according to visual field location and visual category ^5,72–74^. In fact, the face pop out effect ^75,76^ seems to point to enhanced top down influences for faces at peripheral locations relative to other visual categories (but see ^77^). Lastly, despite the fact that our results seem to be in agreement with neuroimaging results, it could very well be that the relationship between brain function and behaviour is much more complex and that our results do not reflect a one to one correspondence with functional activity in high level visual cortex. Hence the effects of eccentricity on face and house discrimination could be partially due to the demands imposed on the system by low-level aspects or by top-down attentional mechanisms rather than being attributed solely to face or house related perceptual mechanisms.

In conclusion, our investigations of face and house discrimination at foveal and parafoveal regions revealed that as expected, performance for all visual categories reduced as a function of eccentricity. Our central to parafoveal investigations of upright and inverted faces and houses allowed us to carry out multiple analyses together suggesting that high level visual processing may be reflected in behavioural performance. Furthermore, our behavioural results support the notion that faces are associated with a foveal bias while houses are less affected by eccentricity.

## Supporting information

Supp. Figures SF1, SF2, SF3, Supp. Tables ST1-5, Supp. Methods, Discussion, and Bibliography

## Acknowledgments

This work was supported by ISF grant no. 1485/18 to SGD. We thank Inbal Ziv, Oren Kadosh and Uri Polat for their assistance, the Computer Vision Laboratory, Faculty of Computer and Information Science, University of Ljubljana, Ljubljana, Slovenia, and the Secondary School Centre, Velenje, Slovenia, for allowing us to use the CVL Face Database (http://www.lrv.fri.uni-lj.si/facedb.html).

## Author Contributions

OK, YB, and SGD designed the experiments. OK ran the experiments, OK and YB analysed the data. OK, YB, and SGD interpreted the results. OK and SGD wrote the paper. All authors designed the revised paper, and have reviewed and approved the final manuscript.

## Competing Interests Statement

All authors (OK, YB, and SGD) declare that they do not have any conflicts of interests.

## Data Availability

No datasets were generated or analysed during the current study.

