## Supplementary material for "Investigating face and house discrimination at foveal to parafoveal locations reveals category-specific characteristics": Supp. Figures SF1, SF2, SF3, Supp. Tables ST1-5, Supp. Methods, Discussion, and Bibliography

**Supplementary Figures**

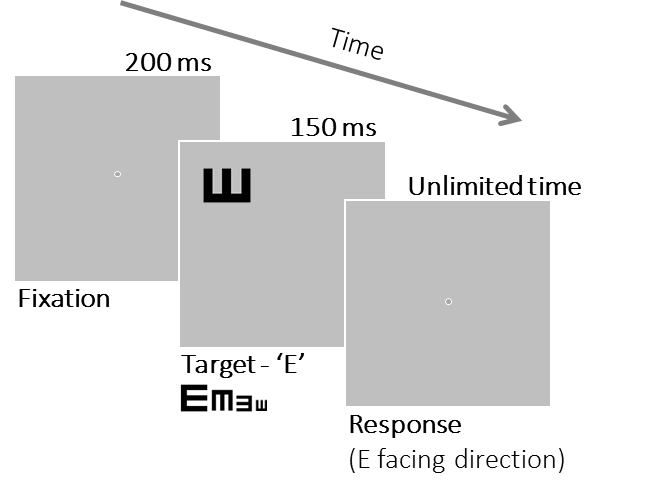

**Supp. Figure SF1**. Timeline of a single trial in the ‘VA’ tumbling E experiment, representing part of a separate staircase procedure performed at 9 different locations from the ones used in the main category discrimination experiments (centre, 2° and 4° eccentricity). Trials from the different staircases (a separate staircase in each location) were interleaved randomly (see “VA tumbling 'E' experiment” section in Methods).

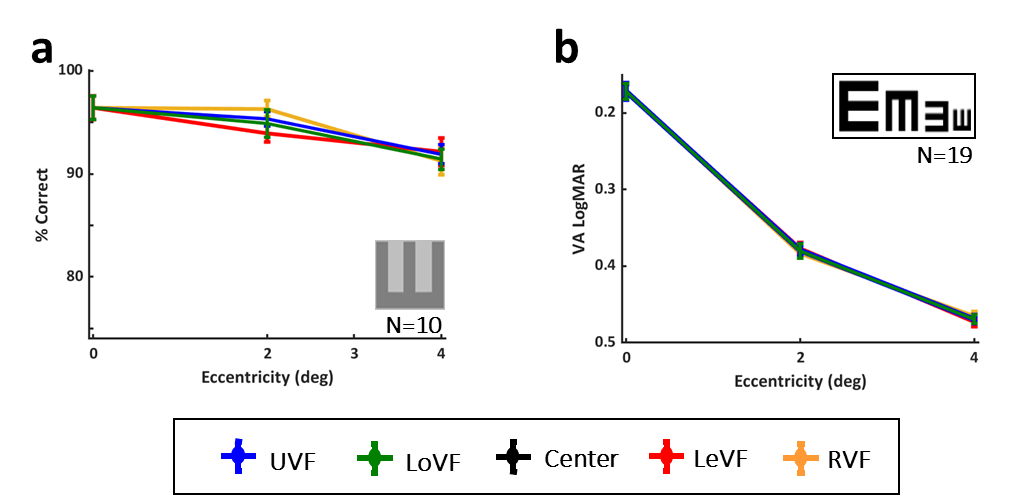

**Supp. Figure SF2**. Accuracy for the E shape discrimination and the VA tumbling E experiments by visual hemifields. Performance for (a) E shape (n=10), (b) tumbling E VA (n=19) declined with eccentricity in each experiment. No significant differences between upper (in blue) and lower (in green) visual fields or between right (in yellow) and left visual (in red) fields were found in these experiments. Error bars represent standard error across participants calculated using the Cousineau method.

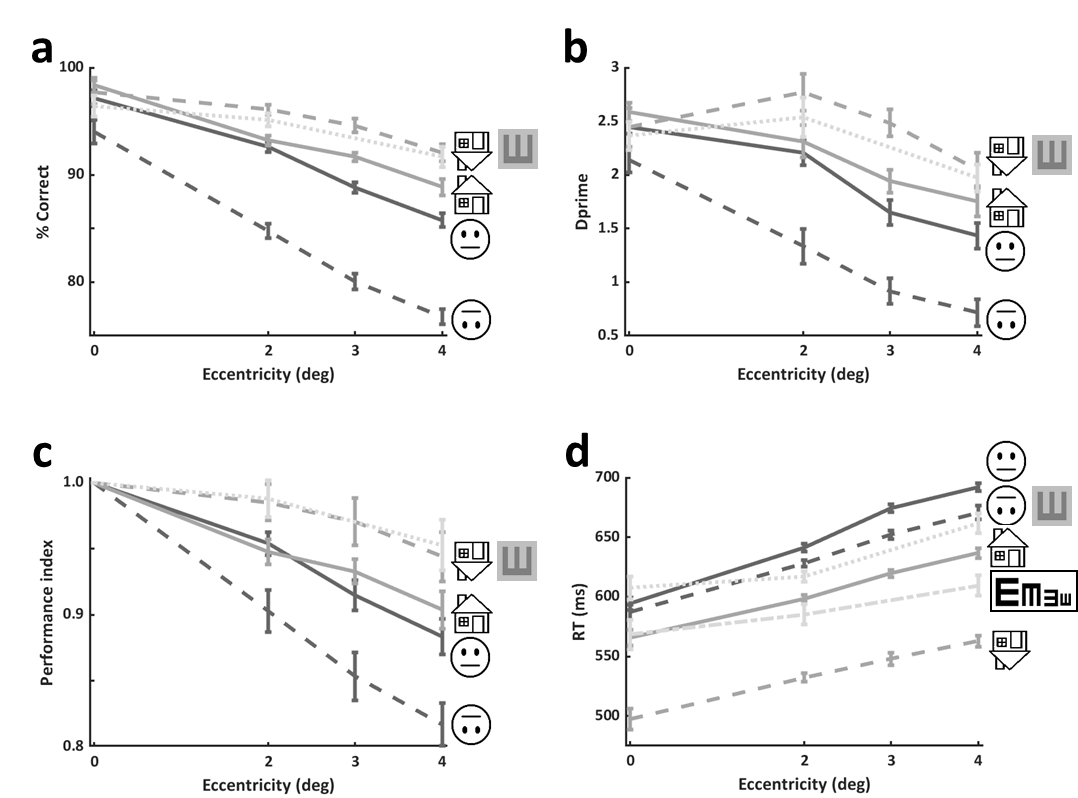

**Supp. Figure SF3**. Joint results across all experiments (all participants) in a similar presentation manner as in Figure 2. Note that all visual categories appear together for presentation purposes only. We refrain from direct between-category comparisons due to the reasons detailed in the main text. (a) Accuracy, (b) dprime, (c) performance index, and (d) RTs; all graphs are based on all study participants (see Methods and Supplementary Table ST1). Error bars in (a), (b), and (d) represent standard errors across participants calculated using the Cousineau method, error bars in (c) represent standard error of the mean (SEM).

**Supplementary Tables**

|  | | **Face discrimination** | | **House discrimination** | | **Shape discrimination** | **VA "Tumbling E"** |
| --- | --- | --- | --- | --- | --- | --- | --- |
|  |  | **Upright** | **Inverted** | **Upright** | **Inverted** |  |  |
|  |  | (n=29) | (n=22) | (n=22) | (n=15) | (n=10) | (n=19) |
| **Accuracy** | Centre | 97.17 ± 0.73 | 94.03 ± 1.10 | 98.34 ± 0.74 | 97.70 ± 1.09 | 96.40 ± 0.95 | 0.17 ± 0.009 |
|  | 2° | 92.63 ± 0.48 | 84.78 ± 0.68 | 93.20 ± 0.47 | 96.09 ± 0.44 | 95.11 ± 0.60 | 0.38 ± 0.006 |
|  | 3° | 88.81 ± 0.49 | 80.06 ± 0.71 | 91.68 ± 0.39 | 94.60 ± 0.66 | NA | NA |
|  | 4° | 85.75 ± 0.66 | 76.77 ± 0.68 | 88.85 ± 0.74 | 92.06 ± 0.78 | 91.67 ± 0.93 | 0.46 ± 0.006 |
| **RT (ms)** | Centre | 594.32 ± 5.18 | 586.95 ± 6.40 | 565.79 ± 6.65 | 497.46 ± 8.71 | 607.50 ± 9.55 | 568.21 ± 12.3 |
|  | 2° | 641.22 ± 3.38 | 628.15 ± 2.87 | 598.40 ± 2.87 | 632.23 ± 3.44 | 617.05 ± 4.30 | 585.26 ± 8.56 |
|  | 3° | 674.46 ± 3.26 | 652.09 ± 3.74 | 619.50 ± 2.99 | 547.83 ± 5.24 | NA | NA |
|  | 4° | 692.06 ± 3.51 | 670.65 ± 5.91 | 636.68 ± 4.25 | 562.80 ± 4.73 | 661.65 ± 8.01 | 609.55 ± 8.49 |
| **Performance index** | Centre | 1 ± 0 | 1 ± 0 | 1 ± 0 | 1 ± 0 | 1 ± 0 | NA |
|  | 2° | 0.95 ± 0.008 | 0.90 ± 0.016 | 0.94 ± 0.009 | 0.98 ± 0.013 | 0.98 ± 0.014 | NA |
|  | 3° | 0.91 ± 0.011 | 0.85 ± 0.018 | 0.93 ± 0.009 | 0.97 ± 0.018 | NA | NA |
|  | 4° | 0.88 ± 0.013 | 0.81 ± 0.016 | 0.90 ± 0.014 | 0.94 ± 0.018 | 0.98 ± 0.019 | NA |
| **Dprime** | Centre | 2.44 ± 0.05 | 2.13 ± 0.11 | 2.58 ± 0.08 | 2.44 ± 0.17 | 2.36 ± 0.13 | NA |
|  | 2° | 2.20 ± 0.11 | 1.33 ± 0.16 | 2.31 ± 0.14 | 2.77 ± 0.17 | 2.53 ± 0.18 | NA |
|  | 3° | 1.64 ± 0.11 | 0.91 ± 0.12 | 1.94 ± 0.10 | 2.48 ± 0.12 | 1.97 ± 0.12 | NA |
|  | 4° | 1.43 ± 0.11 | 0.71 ± 0.12 | 1.75 ± 0.14 | 2.04 ± 0.16 | 1.75 ± 0.14 | NA |

**Supplementary Table ST1**. Summary of experimental results for each experiment by eccentricity. Mean ± SE are provided for accuracy, d-prime, and performance index; median ± SE are provided for reaction times for all experiments. Accuracy (discrimination experiments) is reported in % correct, visual acuity (tumbling E experiment) is reported in LogMAR units. Reaction time (RT) is reported in ms. Performance index represents proportion out of 1. NA - data not available.

|  | | **Face discrimination** | | **House discrimination** | | **Shape discrimination** | **VA "Tumbling E"** |
| --- | --- | --- | --- | --- | --- | --- | --- |
|  |  | **Upright** | **Inverted** | **Upright** | **Inverted** |  |  |
|  |  | (n=29) | (n=22) | (n=22) | (n=15) | (n=10) | (n=19) |
| **Centre** | 0° | 97.12 ± 0.84 | 94.03 ± 1.26 | 98.34 ± 0.81 | 97.70 ± 1.25 | 96.40 ± 1.14 | 0.17 ± 0.010 |
| **UVF** | 2° | 92.99 ± 0.63 | 85.20 ± 1.07 | 92.76 ± 0.55 | 96.32 ± 0.70 | 95.33 ± 0.71 | 0.37 ± 0.008 |
|  | 3° | 88.59 ± 0.83 | 80.72 ± 0.94 | 92.07 ± 0.76 | 94.27 ± 0.94 | NA | NA |
|  | 4° | 86.09 ± 0.89 | 77.98 ± 1.29 | 88.91 ± 0.68 | 92.89 ± 0.79 | 31.91 ± 0.92 | 0.46 ± 0.004 |
| **LoVF** | 2° | 92.27 ± 0.59 | 84.31 ± 0.87 | 93.61 ± 0.57 | 95.88 ± 0.49 | 94.90 ± 1.34 | 0.38 ± 0.009 |
|  | 3° | 89.07 ± 0.81 | 79.52 ± 0.98 | 91.26 ± 0.71 | 94.96 ± 0.75 | NA | NA |
|  | 4° | 85.48 ± 0.88 | 75.78 ± 1.23 | 88.82 ± 0.97 | 91.20 ± 0.90 | 91.41 ± 0.99 | 0.46 ± 0.005 |
| **RVF** | 2° | 92.17 ± 0.62 | 84.65 ± 0.94 | 93.74 ± 0.55 | 95.87 ± 0.77 | 96.30 ± 0.78 | 0.38 ± 0.005 |
|  | 3° | 88.36 ± 0.82 | 78.44 ± 1.04 | 91.76 ± 0.59 | 95.01 ± 0.62 | NA | NA |
|  | 4° | 84.73 ± 1.00 | 75.91 ± 1.02 | 88.50 ± 0.76 | 90.88 ± 0.81 | 91.27 ± 1.37 | 0.46 ± 0.006 |
| **LeVF** | 2° | 93.09 ± 0.78 | 84.83 ± 1.03 | 92.67 ± 0.73 | 96.25 ± 0.61 | 93.31 ± 0.84 | 0.37 ± 0.007 |
|  | 3° | 89.21 ± 0.58 | 81.67 ± 0.822 | 91.67 ± 0.53 | 94.27 ± 0.89 | NA | NA |
|  | 4° | 86.80 ± 0.75 | 77.49 ± 0.85 | 89.17 ± 0.86 | 93.19 ± 0.83 | 92.15 ± 1.35 | 0.47 ± 0.005 |

**Supplementary Table ST2**. Summary of experimental accuracy results for each experiment by visual hemifield. Notations as in Supp. Table ST1.

|  |  | **Factor** | **Main effect** | **Interaction** |
| --- | --- | --- | --- | --- |
| **Upright face**  **(n=29)** | UVF-LoVF | VF (UVF-LoVF) | F(1,28)=0.124 p=0.727 | F(2,56)=0.399 p=0.673 |
|  |  | Ecc (2°,3°,4°) | F(2,56)=31.746 **p<0.0001** |  |
|  | RVF-LeVF | VF (RVF-LeVF) | F(1,28)=2.694 p=0.111 | F(2,56)=0.439 p=0.646 |
|  |  | Ecc (2°,3°,4°) | F(2,56)=31.547 **p<0.0001** |  |
| **Inverted face**  **(n=22)** | UVF-LoVF | VF (UVF-LoVF) | F(1,21)=1.245 p=0.277 | F(2,42)=0.213 p=0.808 |
|  |  | Ecc (2°,3°,4°) | F(2,42)=30.540 **p<0.0001** |  |
|  | RVF-LeVF | VF (RVF-LeVF) | F(1,21)=3.691 p=0.068 | F(2,42)=1.129 p=0.332 |
|  |  | Ecc (2°,3°,4°) | F(2,42)=31.670 **p<0.0001** |  |
| **Upright house**  **(n=22)** | UVF-LoVF | VF (UVF-LoVF) | F(1,21)=0.0003 p=0.984 | F(2,42)=0.742 p=0.482 |
|  |  | Ecc (2°,3°,4°) | F(2,42)=12.535 **p<0.0001** |  |
|  | RVF-LeVF | VF (RVF-LeVF) | F(1,21)=0.109 p=0.744 | F(2,42)=0.948 p=0.395 |
|  |  | Ecc (2°,3°,4°) | F(2,42)=12.728 **p<0.0001** |  |
| **Inverted house**  **(n=15)** | UVF-LoVF | VF (UVF-LoVF) | F(1,14)=0.762 p=0.397 | F(2,28)=1.107 p=0.344 |
|  |  | Ecc (2°,3°,4°) | F(2,28)=9.837 **p=0.0006** |  |
|  | RVF-LeVF | VF (RVF-LeVF) | F(1,14)=0.917 p=0.354 | F(2,28)=2.761 p=0.080 |
|  |  | Ecc (2°,3°,4°) | F(2,28)=9.730 **p=0.0006** |  |
| **Shape**  **(n=10)** | UVF-LoVF | VF (UVF-LoVF) | F(1,9)=0.225 p=0.646 | F(1,9)=0.001 p=0.978 |
|  |  | Ecc (2°,4°) | F(1,9)=7.859 **p=0.020** |  |
|  | RVF-LeVF | VF (RVF-LeVF) | F(1,9)=0.529 p=0.485 | F(1,9)=1.252 p=0.292 |
|  |  | Ecc (2°,4°) | F(1,9)=7.610 **p=0.022** |  |
| **VA**  **"Tumbling E"**  **(n=19)** | UVF-LoVF | VF (UVF-LoVF) | F(1,18)=0.002 p=0.964 | F(1,18)=0.006 p=0.938 |
|  |  | Ecc (2°,4°) | F(1,18)=156.821 **p<0.0001** |  |
|  | RVF-LeVF | VF (RVF-LeVF) | F(1,18)=0.015 p=0.902 | F(1,18)=5.405 **p=0.032** |
|  |  | Ecc (2°,4°) | F(1,18)=156.821 **p<0.0001** |  |

**Supplementary Table ST3**. Summary of statistical visual hemifield analyses across experiments. Statistical analyses were performed using repeated measures two-way ANOVA with eccentricity and visual hemifield on accuracy (or VA results) for each experiment. This was followed by Bonferroni/Dunn post-hoc tests. F statistics are provided with corresponding p values. Values in bold represent significant results. Note that since none of the post-hoc visual hemifield analyses survived Bonferroni/Dunn correction, these are not included in the table).

|  | | Upright houses | | Faces |
| --- | --- | --- | --- | --- |
|  |  | Version 1 | Version 2 |  |
| **Physical difference** | | 23.63 ± 0.57 | 17.78 ± 0.66 | 10.44 ± 0.42 |
| **Accuracy** | Centre | 98.97 ± 0.82 | 97.60 ± 0.91 | 97.17 ± 0.73 |
|  | 2° | 93.60 ± 0.89 | 92.48 ± 0.57 | 92.63 ± 0.48 |
|  | 3° | 92.67 ± 0.68 | 90.79 ± 0.48 | 88.81 ± 0.49 |
|  | 4° | 91.87 ± 0.74 | 87.05 ± 1.06 | 85.75 ± 0.66 |
| **RT** | Centre | 571 ± 7.26 | 566.15 ± 7.39 | 594.32 ± 5.18 |
|  | 2° | 605.54 ± 10.59 | 600.13 ± 4.29 | 641.22 ± 3.38 |
|  | 3° | 623.38 ± 6.59 | 621.15 ± 2.66 | 674.46 ± 3.26 |
|  | 4° | 643.59 ± 8.22 | 641.29 ± 4.63 | 692.06 ± 3.51 |

**Supplementary Table ST4**. Summary of physical differences and experimental results according to experimental version (upright faces (n=29), upright houses Version 1 (n=22) or Version 2 (n=22)). Accuracy and RT are provided by eccentricity. Conventions as in Supp. Table ST1.

|  | **Factor** | **Main effect** | **Interaction** | **Post hoc (Bonferroni/Dunn)** |
| --- | --- | --- | --- | --- |
| Physical difference | Version | F(2,72)=143.421 **p<0.0001** |  |  |
|  | Version1, Version2 |  |  | **p<0.0001** |
|  | Version2, Faces |  |  | **p<0.0001** |
| Accuracy | Version | F(2,280)=9.234 **p=0.0001** | F(6,280)=1.296 p=0.259 |  |
|  | Version1, Version2 |  |  | **p=0.0044** |
|  | Version2, Faces |  |  | p=0.241 |
|  | Ecc | F(3,280)=42.980 **p<0.0001** |  |  |
|  | Center, 2Deg |  |  | **p<0.0001** |
|  | 2Deg, 3Deg |  |  | p=0.0092 |
|  | 3Deg, 4Deg |  |  | **p=0.003** |
| RT | Version | F(2,280)=6.791 **p=0.001** | F(6,280)=0.145 p=0.989 |  |
|  | Version1, Version2 |  |  | p=0.795 |
|  | Version2, Faces |  |  | **p=0.001** |
|  | Ecc | F(3,280)=10.016 **p<0.0001** |  |  |
|  | Center, 2Deg |  |  | p=0.012 |
|  | Center, 3Deg |  |  | **p<0.0001** |
|  | 3Deg, 4Deg |  |  | p=0.221 |

**Supplementary Table ST5**. Physical difference statistical analyses. Statistical analyses to compare the physical differences between upright houses version 1 (n=22), version 2 (n=22) and upright faces (n=29), and relate them to performance were performed using non-repeated measures 1-way ANOVA (physical difference) or 2-way ANOVA (accuracy and RT, all participants). These were followed by Bonferroni/Dunn post-hoc tests when a main effect was found. F statistics are provided with corresponding p values. Values in bold represent significant results.

**Supplementary Methods**

**Shape discrimination experiment**

The experimental design was identical to that used in the face discrimination paradigm except the first and second stimulus in each trial were a gray E on a gray background facing one of 2 directions (up or down) randomly. The participants’ task was to determine whether the two consequently presented E's faced the same direction or not. The E stimuli were black on a gray background and subtended 0.38° x 0.38° (width x height). The size of the E’s was determined after preliminary psychophysical assessments revealed that larger E’s (sized similarly to the face or house images) led to ceiling performance at all eccentricities.

**Supplementary Discussion**

Since face-related regions are contiguous to UVF retinotopic representations^1,2^ we hypothesized that face perception ability may show an UVF bias relative to the LoVF. We employed a face discrimination task where a central study face was compared to peripheral (up to 4° in either UVF or LoVF) target face. However, we did not find any such difference between UVF and LoVF performance for the upright faces, or for the other control categories. One possible explanation is that if visual field differences exist for upright faces, they would be evident in more peripheral locations in the VF (> 4°). Another possibility is that VF differences exist for specific perceptual face tasks but not for the face discrimination task we employed.

We also hypothesized that we may find LeVF vs RVF differences for upright faces, given that several earlier studies reported finding a LeVF preference for upright faces ^3–9^. However, we did not find a LeVF vs. RVF difference for any of the visual categories, similar to an earlier study^10^ . As shown by Maurer and Lewis ^11^, at earlier stages of development, visual inputs from each hemifield (R/Le) cross over to the contralateral visual retinotopic cortex. Thus, at early age (e.g. before 4 months of age), LeVF inputs probably predominate in right visual cortex processing ^11^. These may strengthen the reliance of right hemisphere face processing on LeVF inputs and strengthen the connections between them. This is also in line with findings suggesting right hemisphere dominance for face perception ^3,4,7,12–16^. We cannot rule out the possibility that attention may have influenced the results we obtained. Siman Tov et al. (2007)^17^ show that faces appearing in the LeVF activate the contralateral retinotopic cortex and the fronto-parietal attention network to a much greater extent than those appearing in the RVF ^17^. This may indicate that attention is not uniformly distributed and therefore inputs from the LeVF are more prominent to face processing and to our perception. However, since in our experimental paradigm stimuli could randomly appear in any quadrant including central vision, attention may have been spread uniformly across the parafovea and therefore may have modulated any innate bias if one exists.
